# Protective immunity of the primary SARS-CoV-2 infection reduces disease severity post re-infection with Delta variants in Syrian hamsters

**DOI:** 10.1101/2021.11.28.470293

**Authors:** Sreelekshmy Mohandas, Pragya D. Yadav, Anita Shete, Dimpal Nyayanit, Rajlaxmi Jain, Gajanan Sapkal, Chandrasekhar Mote

## Abstract

Delta variant has evolved to become dominant SARS-CoV-2 lineage worldwide and there are reports of secondary infections with varying severity in vaccinated and unvaccinated naturally recovered COVID-19 patients. As the protective immunity following the infection wanes within few months, studies of re-infection after prolonged duration is needed. Hence we assessed the potential of re-infection by Delta, Delta AY.1 and B.1 in COVID-19 recovered hamsters after 3 months of infection. Re-infection with Delta and B.1 variants in hamsters showed reduced viral shedding, lung pathology and lung viral load, whereas the upper respiratory tract viral load remained similar to that of first infection. The reduction in viral load and lung pathology after re-infection with Delta AY.1 variant was not marked. Further we assessed the disease characteristics of Delta AY.1 to understand whether it has any replication advantage over Delta variant and B.1 variant, an early isolate in Syrian hamsters. Body weight changes, viral load in respiratory organs, lung pathology, cytokine response and neutralizing antibody response were assessed. Delta AY.1 variant produced milder disease in comparison to Delta variant and the neutralizing response was similar against Delta, B.1 and B.1.351 variant in contrast to Delta or B.1 infected hamsters which showed a significant reduction in neutralization titres against B.1.351. Elevation of IL-6 levels was observed post infection in hamsters after primary infection. The prior infection could not produce sterilizing immunity but the protective effect was evident following reinfection. This indicates the importance of the transmission prevention efforts even after achieving herd immunity.

**Research in context:** *Evidence before this study:* Secondary infections with Delta variant are being widely reported and there are reports of increased disease severity. Delta sub lineages with K417N substitution has caused concern worldwide due to the presence of the same substitution in Beta variant, a Variant of Concern known for its immune evasion. The information on the biological characteristics of this sub lineage is also scanty.

*Added value of this study:* The present study showed that the secondary infection with Delta variant does not show any evidence of increased disease severity in hamster model. Delta AY. 1 variant produces mild disease in Syrian hamsters in contrast to severe disease caused by Delta variant. Delta, B.1 and AY.1 variant infected hamster sera showed comparable cross neutralizing response against each other. In contrast to the lower neutralizing response shown by B.1 and Delta variant infected animals against B.1.351 variant, Delta AY.1 showed comparable response as that with other variants.

*Implications of the available evidence:* SARS-CoV-2 infections do not produce sterilizing immunity but protect from developing severe disease in case of Delta variant re-infection indicating the importance of the transmission prevention efforts even after achieving herd immunity. Delta AY. 1 infection in hamsters did not show any evidence of speculated immune evasion.

## Introduction

B.1.617.2 lineage of SARS-CoV-2 was first detected in India on 22^nd^ September 2020.^1^ The variant was later categorized as a Variant of Concern and was named as Delta variant by World Health Organization on 31 May 2021.^2^ The variant has spread at an alarming rate to become the most dominant SARS-CoV-2 lineage circulating globally and has spread to 185 countries as on 21^st^ September 2021.^2^ The amino acid substitutions in the spike protein of Delta variant like D614G, T478K, P681R and L452R are known to affect transmissibility and neutralization.^2^ The variant was responsible for the rise in COVID-19 cases in 2021 in many countries like India, United Kingdom, Fiji, South Africa, in parts of Asia, the United States, Australia, and New Zealand.^3^ B.1.617.2 variant have been further subdivided into variants from Delta AY.1 to AY.33 in the Pango lineage designation system.^1^ Among these sub lineages, AY.1 and AY.2 possess K417N substitution which is also present in B.1.351 variant speculating its ability for immune evasion.^1^ As of 30 September 2021, AY.1 lineage has been detected in at least 44 countries and AY.2 in 8 countries.^3^ AY.1 has amino acid substitutions at T19R, E156G, 157/158 del, W258L, K417N, L452R, T478K, D614G D950N and P681R.^1^ The information on the biological characteristics of these sub lineages including transmissibility, disease severity and immune evasion are still unknown.

SARS-CoV-2 generates neutralising antibody response after infection in humans, but protective immune titre to prevent a subsequent infection is not yet understood.^4^ In case of other human coronaviruses (HCoV), waning of immunity is observed in 1 to 3 years and reinfection events have been reported as a common feature of HCoV-NL63, HCoV-229E, HCoV-OC43 and HCoV-HKU1.^5,6^ After the natural SARS-CoV-2 human infection, immune response is suspected to persist for about 90 days in most patients.^4^ SARS-CoV-2 re-infection cases with varied disease severity have been published from many countries.^7,8,9,10,11^ The speculated reasons for re-infection are infection with a higher virus dose/another virulent strain, antibody-dependent enhancement and waning of immune response.^8^ Laboratory studies have shown that the duration of infection-acquired immunity is inconsistent and the response against variants of concern also differ.^12^ The risk of re-infection also depends on host susceptibility, vaccination status, and exposure to COVID-19 patients during infectious phase.^13^ Understanding the potential risk of a re-infection is important in improving COVID-19 prevention and control measures. Re-infection studies in the population are very less and the rate of re-infection is also not clear.^14^ There are few reports of aggravated disease severity in case of Delta variant re-infection.^15,16^ The impact of the immunity on threat of re-infection posed by different variants still needs to be understood. Such studies are important in making policy decisions which rely on the herd immunity like vaccination.

Animal models are important in understanding virus properties, disease pathogenesis, measuring efficacy of countermeasures etc. Syrian hamsters have been widely used in studying SARS-CoV-2 disease characteristics. Our previous studies have shown the pathogenicity and immune evasive properties of the Delta variant in hamsters.^17^ Few animal model studies also indicate the neutralizing antibody response generated after primary SARS-CoV-2 infection can reduce the viral load and severity of a second infection.^18,19^ The degree of protection against a secondary infection by Delta lineage variants is still not clear.

Here we have studied the re-infection potential of Delta, Delta AY.1 and B.1 in B.1 infection recovered hamsters post 3 months of infection and also the pathogenicity of Delta AY. 1 in comparison with Delta and B.1 in Syrian hamsters.

## Methods

### Ethics statement

The ethical approval for the study was received from Institutional Animal Ethics Committee (Approval no: NIV/IAEC/2021/ MCL/01), ICMR-National Institute of Virology, Pune and all the animal experiments were performed in adherence with the guidelines of the Committee for the Purpose of Control and Supervision of Experiments on Animals, Government of India.

### Virus and cells

SARS-CoV-2 variants B.1 (GISAID accession no: EPL_ISL_825084), Delta (GISAID accession no.: EPI_ISL_2400521) and Delta AY.1 (GISAID accession no. EPI_ISL_2671901) isolated from nasopharyngeal swabs of COVID-19 patients were used for the study. The isolates were passaged twice in Vero CCL81 cells and titrated to measure the 50% tissue culture infective dose (TCID50) as per the Reed and Muench method. The variants used in the study had the following amino acid substitutions in the spike protein. Delta AY.1 variant had D614G, E156G, F157del, K417N, L452R, P681R, R158del, T19R, T95I and T478K substitutions, Delta variant had A222V, D614G, D950N, G142D, L452R, P681R, T19R, T478K and B.1 variant had D614G substitution in the spike protein.

### Animal experiments

The experiments were performed in the Containment Facility of ICMR-National Institute of Virology, Pune. For the re-infection study, 12 female hamsters, 16-18 weeks old which were previously infected with B.1 variant of SARS-CoV-2 (with an infectious dose of 10^4,5^ TCID50) after 3 months of initial infection were used (Figure 1a). IgG response and neutralizing antibody levels were checked and animals were divided into 3 groups randomly.

**Figure 1:**
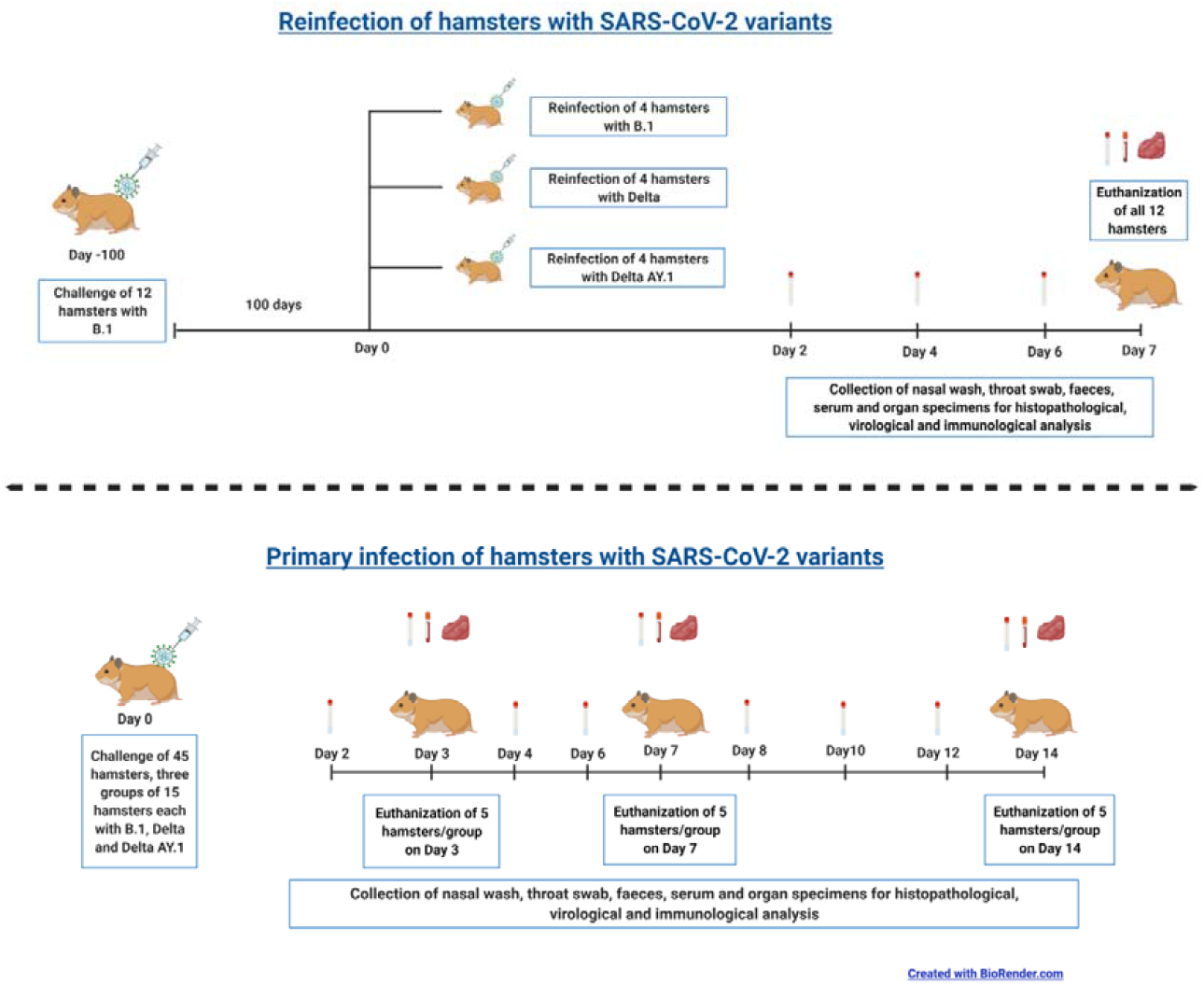
Study design. a) Summary of the Delta AY.1 vs Delta and B. 1 pathogenicity study b) Summary of the re-infection study

The hamsters were divided into 4 animal per group and were re-infected with Delta/Delta AY.1/B.1variants with a virus dose of 10^5^ TCID50. Swab samples were collected on 2, 4, 6 DPI and body weight change was monitored till 7 days. The hamsters were sacrificed on 7DPI to collect organ and blood samples.

As control for the re-infection study and also to understand the disease characteristics of Delta AY.1, three study groups of 17 female, Syrian hamsters of 12-14 week age each were included in the study to assess pathogenicity and a virus dose of 10^5^ TCID50 of Delta/ Delta AY.1/B.1 was used intranasally to inoculate the hamsters (Figure 1b). Swab samples (n=7) were collected on alternate days during the study period. Hamsters were observed for a period of 14 days for body weight loss and 5 hamsters/group were sacrificed on 3, 7- and 14-days post infection (DPI) to collect organs and blood samples.

### Viral load estimation

Nasal wash, throat swab and organ tissue samples were used for viral load estimation. Organ samples collected during necropsy were weighed and homogenized in sterile media using beads in a tissue lyser machine (Qiagen, Germany). The lysate was used for RNA extraction using the MagMAX™ Viral/Pathogen Nucleic Acid Isolation Kit as per the manufacturer’s instructions. Quantitative real-time RT-PCR was performed for the E gene of SARS-CoV-2 using published primers to estimate the genomic viral RNA load and for the N gene of SARS-CoV-2 using published primers to estimate the subgenomic viral RNA load.^20,21^

### Anti-SARS-CoV-2 IgG detection

The serum samples were tested for IgG antibodies by an in-house developed ELISA^22^. Briefly, inactivated SARS-CoV-2 antigen/ Vero CCL81 cell lysate coated microtiter plates were blocked with liquid plate sealer. Hamster sera samples diluted 1: 100 were added and incubated for 60 minutes at 37°C. The plates were washed following incubation and 1:3000 dilution of anti-hamster IgG-horse radish peroxidase (Thermoscientific, USA) was added and incubated for 60 minutes. The plates were washed and substrate was added to each well for color development. The reaction was terminated with sulfuric acid and the absorbance was measured at 450 nm using an ELISA reader. The assay was performed in duplicate and the assay cut off was set at an optical density value of 0.2 and positive/negative ratio of 1.5.

### Serum neutralizing antibody level estimation

Plaque Reduction Neutralization test (PRNT) was performed against B.1, Delta, Delta AY.1 and Beta (B. 1.351) variants as described earlier^23^. Diluted sera were mixed with virus containing a 50-60 plaque forming units/0.1 ml and the virus-sera mixture was incubated for 60 minutes and added in a tissue culture plate with Vero CCL-81 monolayer. After 60 minutes, the mixture was aspirated and media with 2% carboxymethyl cellulose with 2% fetal bovine serum was added. After an incubation period of 4 days, the media was decanted and amido black staining was performed. The plaques were counted and PRNT50 titers were calculated.

### Serum cytokine level estimation

ELISA based estimation (Immunotag, USA) was performed to assess the levels of IL-4, IL-6, IL-10, IFN-γ and TNF-α in hamster sera samples as per the manufacturer’s instructions.

### Lung histopathological evaluation

Formalin fixed lung tissue samples were processed using an automated tissue processor and were stained by routine hematoxylin and eosin staining. The samples were coded and were blindly scored. The bronchiolar (degeneration, epithelial loss), alveolar parenchymal (edema, exudation, mononuclear infiltration, emphysema, pneumocyte hyperplasia, septal thickening) and vascular lesions (congestion, haemorrhages, perivascular infiltrations) were graded for severity on a score from 0 to 4.

### Data analysis

Graph pad Prism version 9.2.0 software was used for the descriptive statistics and statistical analysis. Non parametric Mann Whitney tests were used for the analysis. The p-values less than 0.05 were considered statistically significant.

## Results

### Body weight changes and immune response post Delta AY.1/Delta/B.1variant infection

A mean body weight loss of −9.3 ± 6.5 (mean ± standard deviation) % was observed in Delta infected group (*p=0.0248 vs* Delta AY.1 and *p=0.0082 vs* B.1) on 6 DPI whereas the Delta AY.1 and B.1 infected group animals showed a average weight gain of 0.23 ± 5.18 % and 1.4 ± 3.34 % respectively (Fig. 2a). Anti-SARS CoV-2 IgG response could be observed from day 3 in all infected groups with an mean optical density (OD) ± standard deviation (SD) of 0.65± 0.45, 1.00 ± 0.67, 1.02 ± 0.40 on 3DPI and 1.25 ± 0.55, 0.62± 0.13 and 0.62 ± 0.11 on 14 DPI for Delta AY.1, Delta and B.1 variant respectively (Fig. 2b).

**Figure 2:**
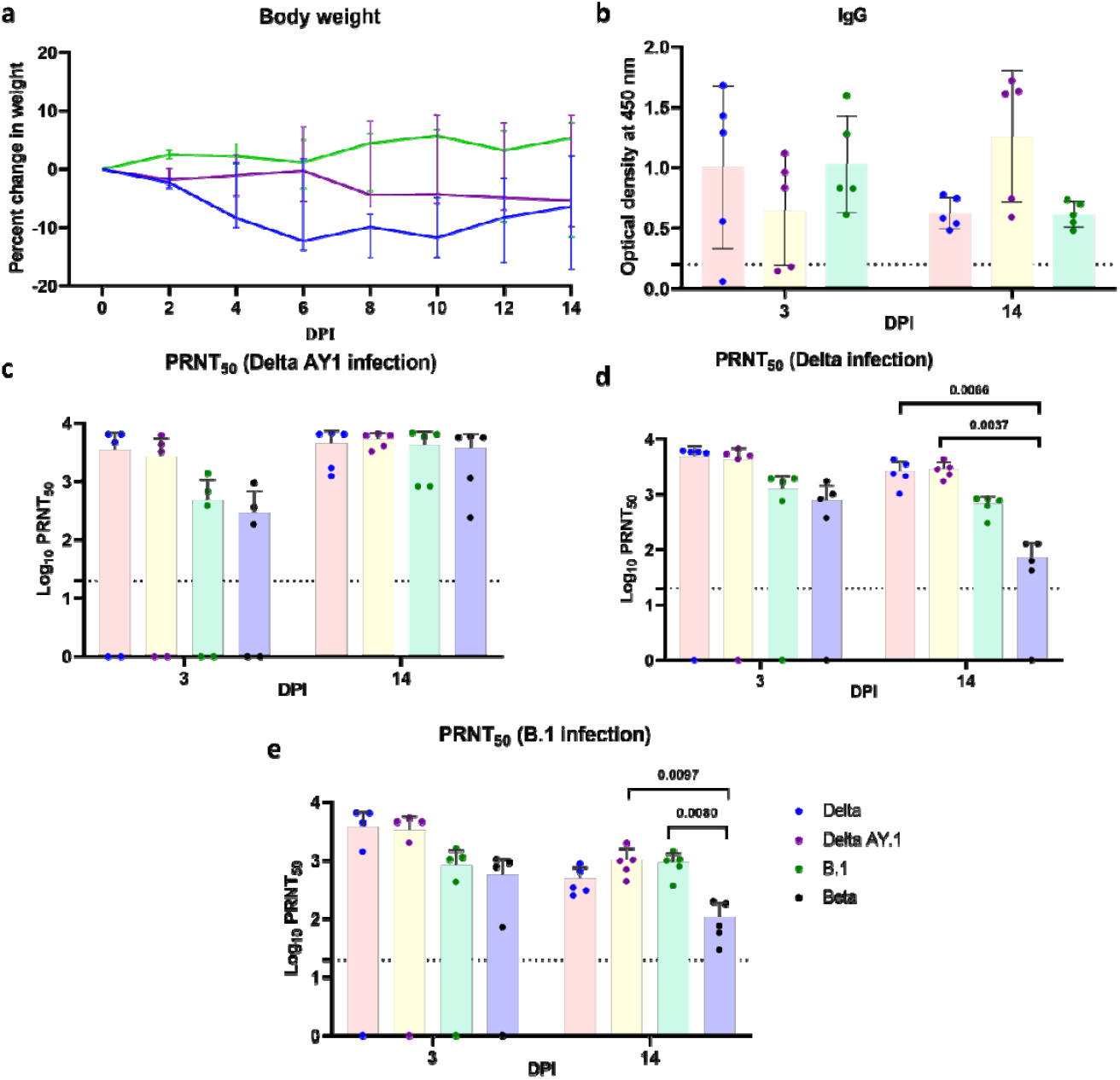
Body weight loss and immune response in hamsters post infection. **a)** Line graph depicting the body weight loss in hamsters post first infection. **b)** Scatter plot depicting IgG response in hamsters post first infection (n=5). Scatter plot depicting **c)** PRNT50 titres against variants in Delta AY.1 infected hamsters (n=5) **d)** PRNT50 titres against variants in Delta infected hamsters, [*p=0.0066 (Delta vs Beta), Mann Whitney test*, n=5] and **e)** PRNT50 titres against variants in B.1 infected hamsters *[p=0.0080 B.1 vs Beta, Mann Whitney test, n=5)*. Limit of detection is assay is depicted as the dotted line.

Neutralizing antibodies could also be detected in the animals from 3DPI. The mean ± SD of neutralizing antibody titre on 14 DPI in Delta AY.1 infected group against Delta AY.1, Delta, B.1 and B.1.351 variant were 5265±1504, 4544±2824, 4159±3062 and 3735±2804 respectively. In case of Delta and B.1 infected groups, mean neutralizing antibody response against Delta AY.1, Delta, B.1 and B.1.351 variant were 2848 ±978, 2667 ± 1275, 680 ±234, 73 ± 57 and 1020 ± 59, 485 ± 269, 925 ±398 and 109 ± 77 respectively. Against B.1.351, a significant reduction in neutralizing antibody titre was observed with Delta *(p= 0.0066)* and B.1 *(p= 0.0080)* variant infected animal sera whereas comparable response was observed in case of AY.1 infected group (Fig.2c–2e).

Among the cytokines analysed in serum, IL-4, IL-10, Interferon-gamma and TNF-alpha showed no significance in comparison with control animal sera on 3, 7 and 14 DPI, whereas IL-6 levels were found higher in infected groups irrespective of variant of infection. B.1 group hamsters showed a significantly higher IL-6 level on 3 (p=0.0037), 7(p=0.0011) and 14 (p=0.0327) DPI and Delta group on 14 DPI (p=0.0277) (Fig. 3a-e).

**Figure 3:**
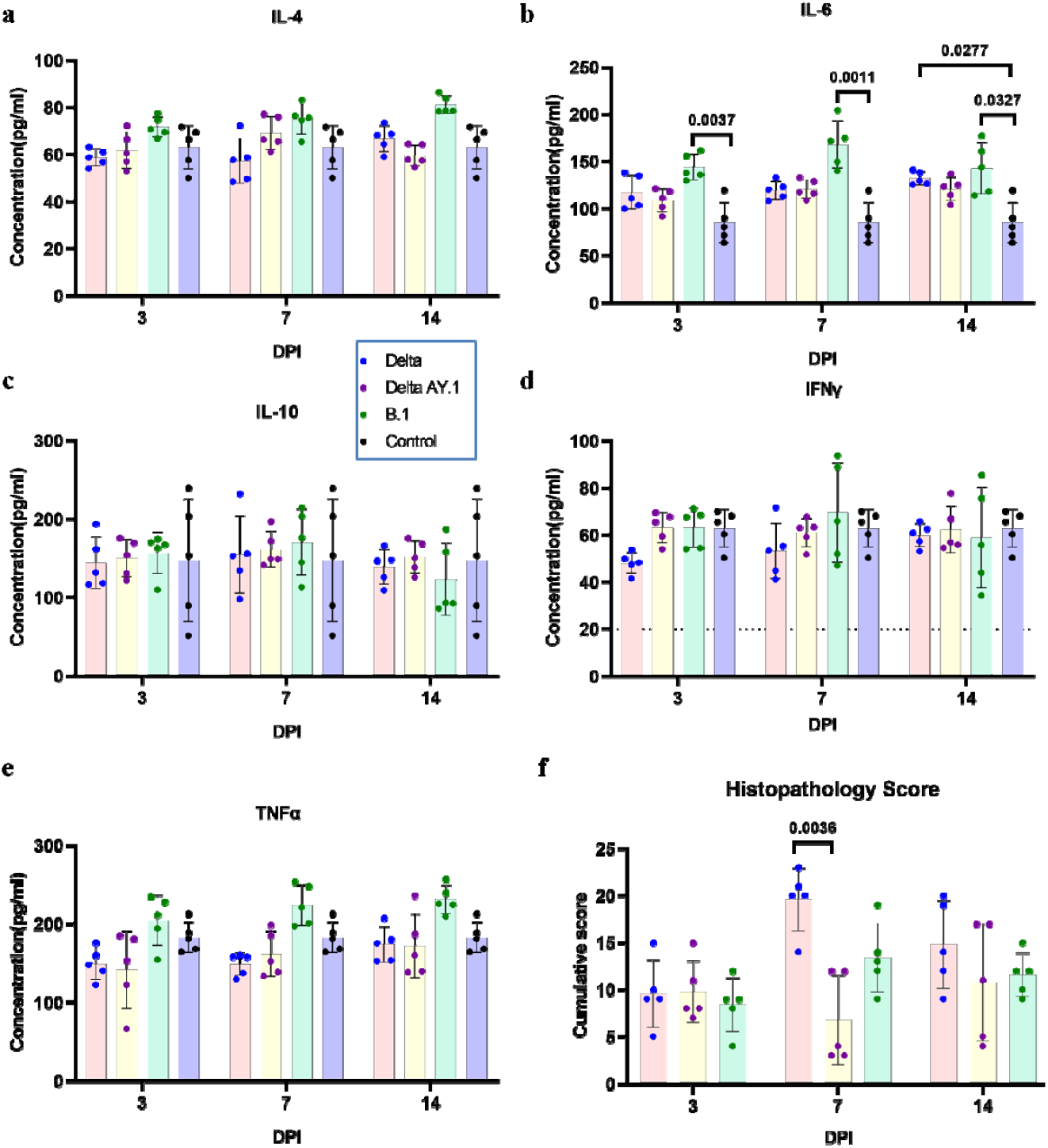
Serum cytokine levels and the histopathology score in hamsters post SARS CoV-2 infection. Scatter plot depicting a) IL-4, b) IL-6 *(p=0.0037on 3DPI,p=0.0011 on 7 DPI and p=0.0327 on 14 DPI, B.1 vs Control, p= 0.0277 on 14 DPI, Delta vs Control, Mann Whitney test, n=5)*, c) IL-10, d) IFN-gamma e) TNF-alpha levels in serum of hamsters on 3, 7 and 14 DPI post infection. f) Scatter plot depicting cumulative lung histopathology score in hamsters post infection *(p=0.0036, Delta vs Delta AY.1, Mann Whitney test, n=5)*.

### Lung pathological changes post Delta AY.1/Delta/B.1 variant infection

Grossly 2/10 animals showed haemorrhagic lesions in case of Delta AY.1 and 7/10 animals lesions of Delta variant sacrificed on 7^th^ and 14^th^ DPI. On 3^rd^ DPI, the vascular pathological changes were minimal in all groups and mild pneumonic changes were observed in the alveolar parenchyma. The pneumonic changes were minimal in the Delta AY. 1 group on 7 DPI, which became more pronounced by 14 DPI in 3/5 animals infected. The highest lung cumulative score was observed in Delta infected group on 7 DPI. By 7DPI, inflammatory changes became severe in Delta group characterized by severe congestion/haemorrhages, alveolar consolidation, loss of bronchial epithelium, septal thickening, pneumocyte hyperplasia, cellular infiltration in the alveolar interstitial space, peri bronchial and perivascular area. In case of B.1 pneumonic changes became more pronounced by 7 DPI (Fig 3f).

### Viral load after Delta AY.1/Delta/B.1 variant infection in Syrian hamsters

In the throat swab and nasal wash samples, gRNA could be detected in a decreasing trend till 12 DPI in Delta AY.1 and B.1 infected groups and till 14DPI in Delta variant infected group. The average viral gRNA levels in throat swab was significantly lower in Delta AY.1 infected group in comparison to other variants i.e., on 2^nd^ DPI, *p= 0.0089 vs* B.1, on 4^th^ DPI, *p= 0.0009 vs* B.1, on 6^th^ DPI, *p= 0.0242 vs* Delta, on 8^th^ DPI, *p=0.002 vs* Delta and on 10^th^ DPI, *p=0.0118 vs* Delta (Fig 4a). Sub genomic RNA levels were comparable in throat swab samples on first week and were found significantly lower in Delta AY.1 infected group on 8DPI *(p=0.0036)* in comparison with Delta variant group (Fig 4b). Nasal wash gRNA load was significantly lower in Delta AY.1 group on 2^nd^ (*p=0.0008 vs* Delta), 4^th^ (*p=0.0374 vs* Delta), 8^th^ (p=0.004 *vs* Delta), 10^th^ (p=0.003 *vs* Delta) and on 12^th^ (*p= 0.0005 vs* B.1) DPI (Fig 4c). In nasal wash, sgRNA levels were also significantly lower on 2^nd^ (*p=0.047 vs* Delta), 4^th^ (*p=0.042 vs* Delta), 8^th^ (*p=0.0035 vs* Delta) and 10^th^ (*p=0.0005 vs* Delta) DPI (Fig 4d). Viral gRNA could be detected in the faeces samples during the first week post infection in all the groups and post 8 DPI detection was only in few animals of Delta variant group (Fig 4e). Delta AY.1 group showed significantly lower gRNA on 2^nd^ (*p=0.037 vs* Delta, *p=0.001 vs* B.1), 4^th^ (*p=0.01 vs* Delta, *p=0.006 vs* B.1) and 8^th^ (*p=0.0004 vs* Delta) DPI. The sgRNA could be detected in faeces only on 2^nd^ (*p=0.0085 vs* B.1) and 4^th^ (*p=0.0317 vs* Delta) DPI which were significantly lower in Delta AY.1 group (Fig 4f).

**Figure 4:**
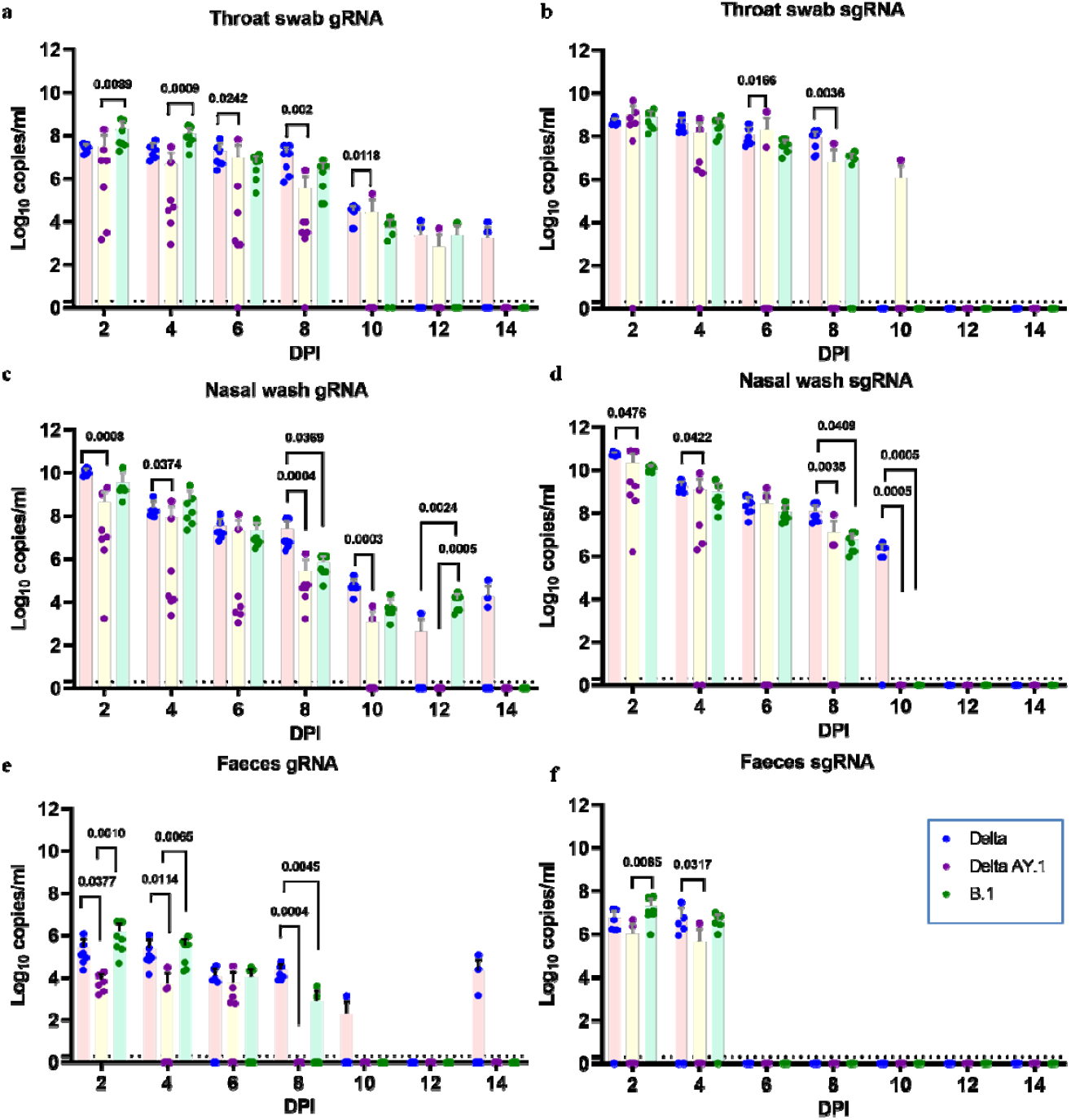
SARS-CoV-2 viral RNA load hamsters after infection. Scatter plot depicting viral gRNA load in hamsters a) throat swab c) nasal wash and e) faeces of hamsters post infection, (*Mann-Whitney test, n=7*). Scatter plot depicting viral sgRNA load in hamsters b) throat swab d) nasal wash and f) faeces of hamsters post infection, *(MannWhitney test, n=7)*.

In nasal turbinates, the gRNA and sgRNA could be detected till 14 DPI in all the groups. Delta AY.1 group did not show any significant difference in the gRNA or sgRNA level of nasal turbinates in comparison to other groups (Fig 5a, 5b). In Delta AY.1 group, a lower mean viral load (mean ± SD = 4.55×10^4^ ± 9.0 ×10^4^) was observed on 7 DPI in lungs, which showed complete clearance of viral gRNA (*p=0.0010 vs* Delta) and sgRNA (*p=0.0057 vs* B.1) by 14 DPI. Other organs like brain (1/5), heart (2/5) and large intestine (2/5) of Delta AY.1 infected group showed sgRNA positivity on 3DPI and none from the Delta and B.1 infected group (Fig.5c,5d).

**Figure 5:**
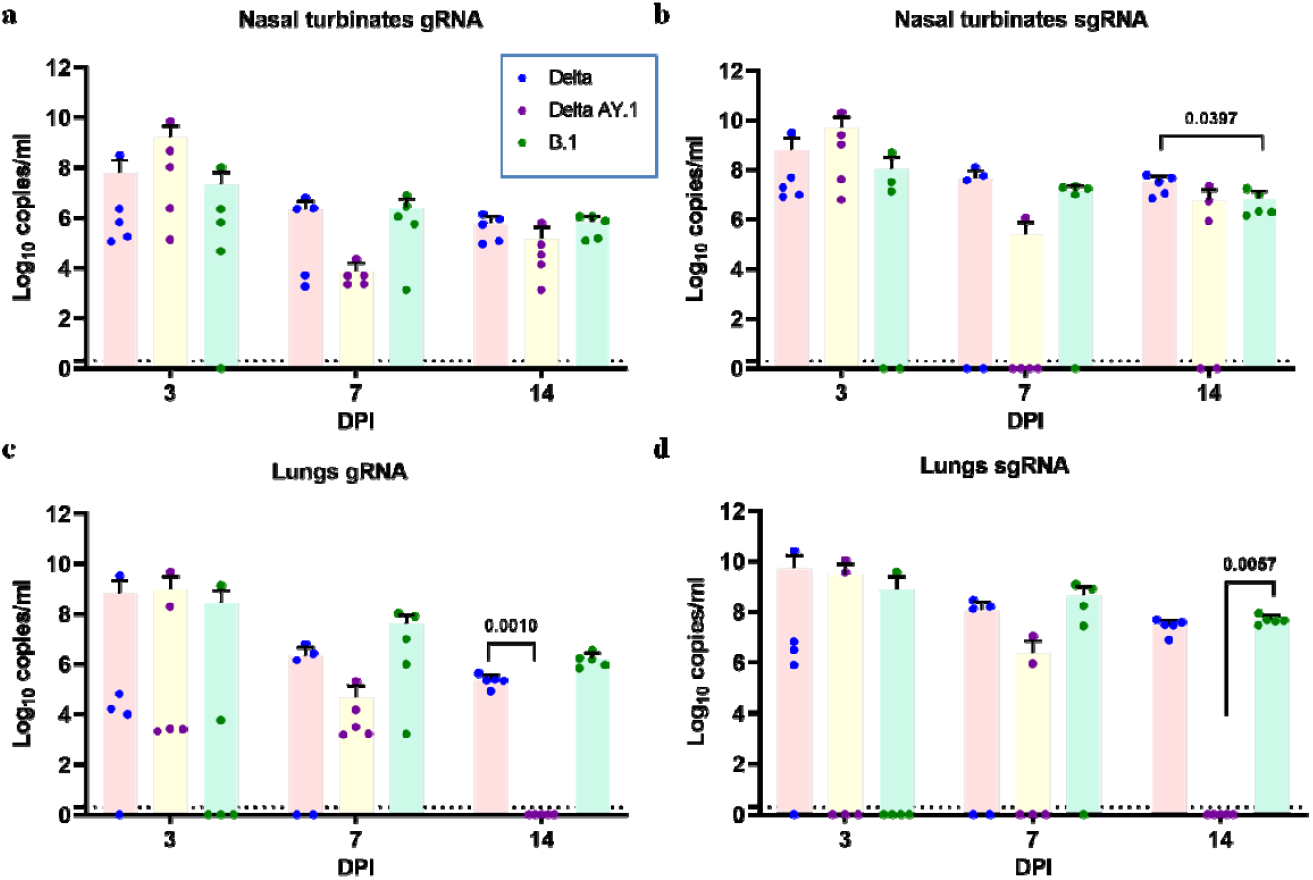
SARS-CoV-2 viral RNA load in organs of hamsters post infection. Scatter plot depicting viral gRNA load in hamsters a) nasal turbinates and c) lungs. Scatter plot depicting viral sgRNA load in b) nasal turbinates *(p=0.0397, Delta vs B.1,Mann Whitney test, n=5)* and d) lungs *(p=0.0057, AY.1 vs B.1,Mann Whitney test, n=5*.

### Delta AY.1, Delta and B.1 variant re-infection in Syrian hamsters

After re-infection, the body weight loss in all the infected groups was minimal irrespective of the variant infected (Supplementary Fig.1a). The mean (±SD) body weight change on 7 DPI post re-infection was 1.74 (± 3.9), 0.67 (±3.8) and −0.3 (±3.05) g in Delta, Delta AY.1 and B.1 groups. The IgG antibody response (mean OD ± SD) measured by ELISA were 0.86 ± 0.45, 0.66±0.18 and 1.03 ± 0.45 on day 0 before re-infection and became 1.54 ± 0.15, 1.40 ±0.14 and 1.54 ±0.07 against Delta, Delta AY.1 and B.1 variant respectively (Supplementary Fig.1b). Post-re infection mean average OD in IgG ELISA on 7 DPI was found increased in all the groups which doubled in case of Delta and B.1 group and tripled in Delta AY.1 group. Post re-infection, the PRNT50 titres also showed rise in titre with all the variants (Table 1). The serum cytokine levels did not show any significant difference between re-infected groups and the uninfected control sera.

**Table 1:** Plaque reduction neutralization titre of hamster sera samples before and after re-infection against SARS-CoV-2 variants.

| Study Group<br>(n=4/<br>group) | PRNT 50 titre against SARS-CoV-2 variants (Mean ± SD) | | | | | | | | |
| --- | --- | --- | --- | --- | --- | --- | --- | --- | --- |
|  | Before re-infection | After re-infection on 7DPI |  |  |  | After primary infection on 7DPI (Control group) |  |  |  |
|  | B.1 | B.1 | Delta | Delta AY.1 | Beta | B.1 | Delta | Delta AY.1 | Beta |
| **B.1** | 1220±818.7 | 6749±0 | 6773±229.3 | 5975±244.74 | 4375±1738.3 | 2342±2565.8 | 3879±2138.2 | 4626±1578.7 | 1542±2934.2 |
| **Delta** | 3675 ± 3155.9 | 6569±207.8 | 6830±116.5 | 6476±348.3 | 4093±2877 | 4327±2386.1 | 5863±1088.9 | 5714±878.5 | 3255±2977.2 |
| **Delta AY.1** | 2213± 1559.1 | 6659±180 | 6773±229.5 | 6888±0 | 5628±1913.9 | 6401±298.4 | 6729±154.0 | 6096±529.9 | 5556±258 |

The viral load in throat swab and nasal wash were significantly lesser in the Delta *(p=0.0286)* and B.1 *(p=0.0286)* re-infected hamsters on 2, 4 and 6 DPI (Fig 6). Even though the viral load in AY.1 group also showed reduction, the values were not statistically significant. The viral load in nasal turbinates were comparable in both re-infected and the control group irrespective of the variant infected (Fig 7). Lungs gRNA level in B.1 infected group was significantly lesser *(p=0.0286)* in B.1 infected group whereas in case of Delta, a slight reduction was seen and AY.1 group showed comparable viral RNA load. In case of primary infection, grossly diffuse haemorrhages in all the lung lobes were seen in 4/5 naïve animals infected with Delta, haemorrhagic foci in one or two lobes in 2/5 animals infected with Delta AY. 1 and few focal haemorrhages in case of B.1 infection (Fig. 8a–8l). In contrary the reinfected animal showed only few focal haemorrhagic foci in case of all variant infections (Fig. 8m–8x). The lungs body weight ratio after the Delta variant re-infection was reduced in comparison to the primary infected animals (Fig.8y). In the re-infected Delta and B.1 group, the lung pathological changes observed on 7DPI were milder in comparison to the naïve infected animals. In case of AY. 1 re-infection, the re-infected group showed similar disease severity as that of 2/5 naive infected animals (Fig 8z).

**Figure 6:**
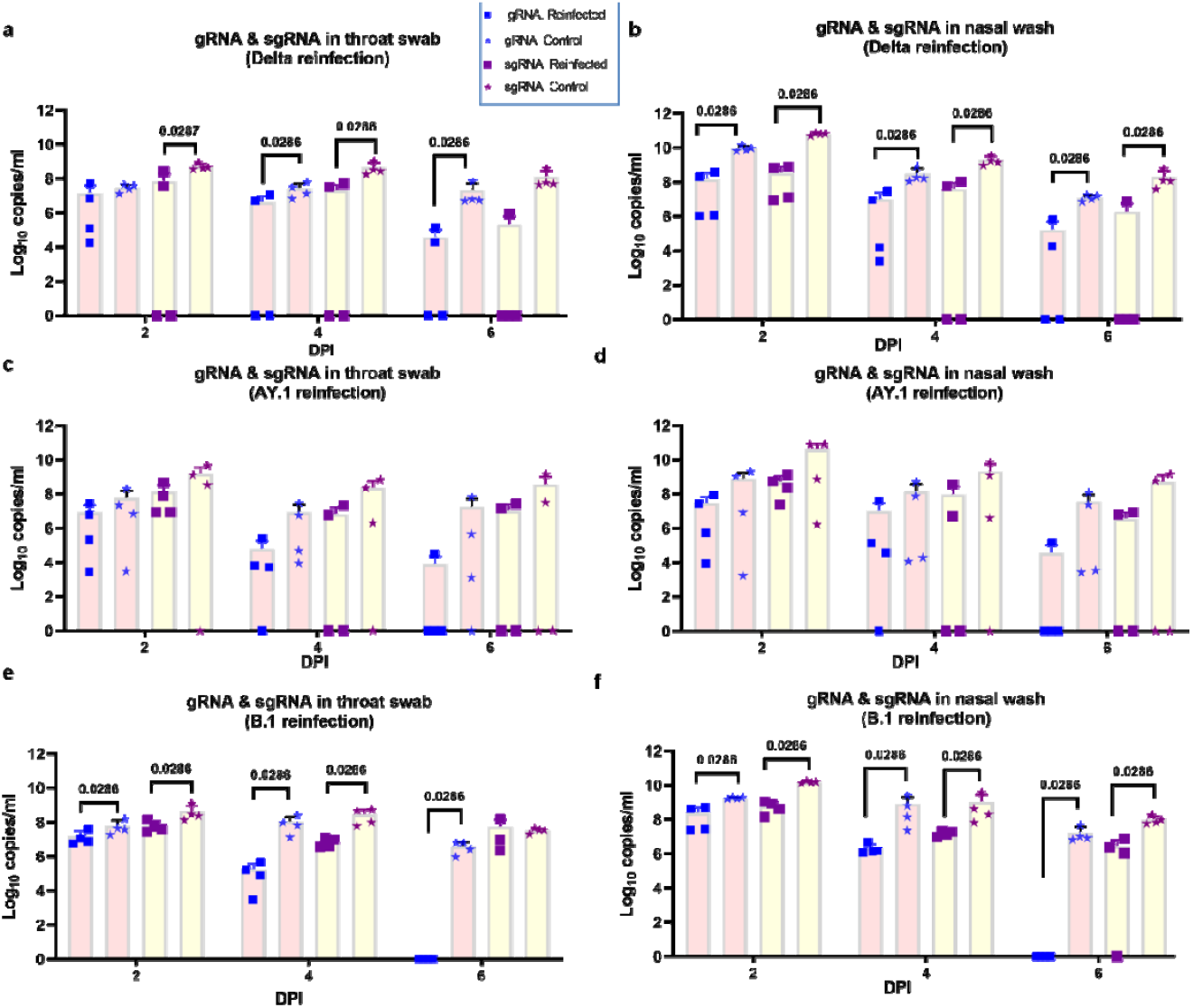
SARS-CoV-2 viral RNA load in hamsters after re-infection. Scatter plot depicting viral gRNA and sgRNA load in a) throat swab, *(p=0.0286, Mann Whitney test, n=4)* and b) nasal wash *(p=0.0286, Mann Whitney test, n=4)* of Delta variant, c) throat swab and d) nasal wash of Delta AY.1 and e) throat swab *(p=0.0286, Mann Whitney test, n=4) and* f) nasal wash *(p=0.0286, Mann Whitney test, n=4)* of B.1 infected hamsters.

**Figure 7:**
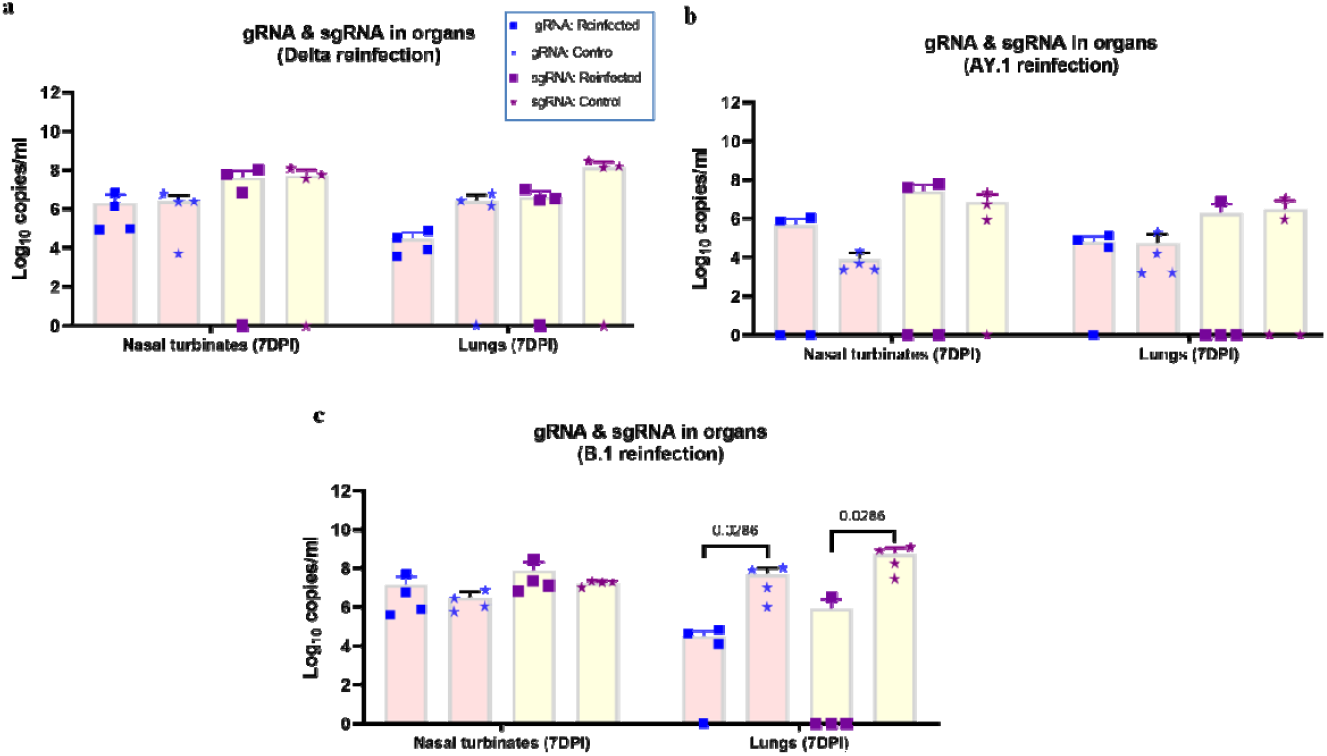
SARS-CoV-2 viral RNA load in organs and lung histopathology score after re-infection. Scatter plot depicting viral gRNA and sgRNA load in nasal turbinates and lungs of a) Delta b) Delta AY.1 and c) B.1 *(p=0.0286, Mann Whitney test, n=4)* infected hamsters.

**Figure 8:**
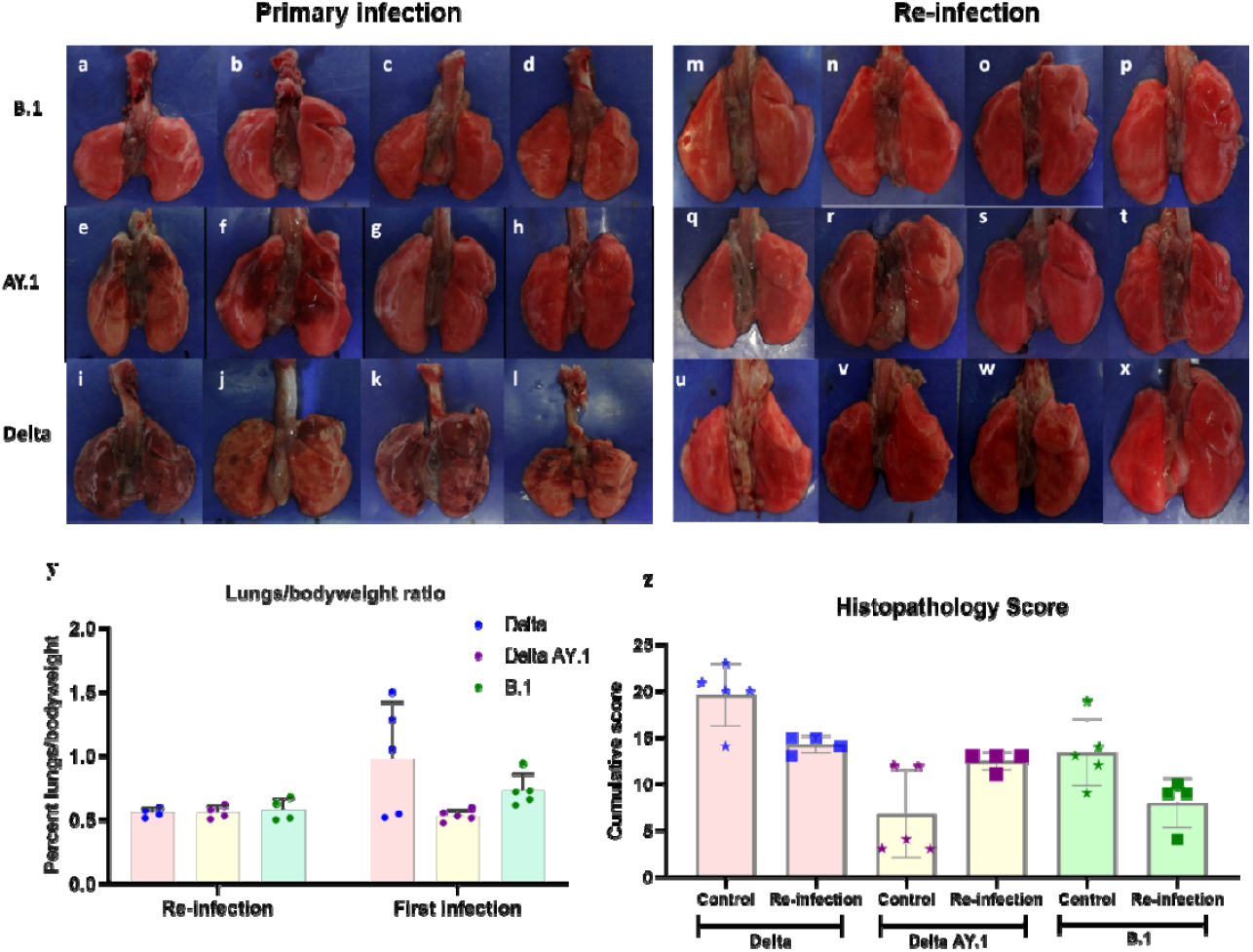
Pathological changes observed in lungs after primary infection and re-infection. Lungs showing (a-d) normal gross appearance following infection with B.1 variant, (e,f) haemorrhages in left and right lower lobes and (g,h) normal gross appearance in Delta AY.1 infected animals and (i-l) diffuse haemorrhages in all lung lobes in Delta infected animals. Lungs showing (m-p) showing normal gross appearance after B.1 re-infection, (q-t) showing normal gross appearance after Delta AY.1 re-infection and (u,x) showing normal gross appearance and (v,w) focal congestion after Delta infection. y) Lungs body weight ratio in hamsters after primary infection and re-infection. z) Scatter plot depicting cumulative lung histopathology score in hamster’s post first infection and re-infection with Delta, Delta AY.1 and B.1 on 7DPI.

## Discussion

Delta variant and its sub lineages have become the dominating SARS-CoV-2 lineage world-wide. Faster transmission, increased disease severity and immune evasion potential of the Delta variant have alerted the scientific community to be vigilant about the mutating variants. ^2,3^ We have studied the potential of re-infection in hamsters by Delta variants 3 months post recovery from SARS-CoV-2 infection, a period till which antibodies are reported to persist in most of the human infections. The disease severity in hamster model depends on the inoculum dose too, as we observed in our earlier study where a lower dose of Delta variant inoculum produced minimal body weight loss and lung disease of moderate severity^17^. Hence we used a higher dose of Delta variant here and observed an increase in the severity of pneumonia and prominent body weight loss and used the same for our re-infection study.

The protective immunity conferred by prior SARS-CoV-2 infection is similar to that of vaccination^23^. Natural infection can generate an effective mucosal immune response unlike intramuscular vaccination.^24^ Neutralizing antibodies could be detected in hamsters 3 months post recovery from infection and the re-infection showed a boosting effect in the neutralization titers. Other than antibody response, cell mediated immunity also play a major role in disease protection^25^. Re-infection with SARS-CoV-2 and other human corona viruses have been reported^6^. The rates of re-infections are common 12 months post initial infection in case of seasonal HCoVs. A trend of reduced virus replication and increase in the neutralizing antibody titre were observed in hamsters following re-infection irrespective of the variant infected. Previous research has shown that prior COVID-19 infection reduces virus replication and thus decrease the transmission efficiency in Syrian hamsters which were reinfected 29 days post first infection.^19^ Even though a reduction was seen in viral shedding, the nasal turbinate viral load remained comparable. The prior infection could not confer sterilizing immunity in the present study also as reported earlier in rhesus macaque and hamster model.^19,26,27^ These studies were performed within a month post recovery from primary infection in contrast to our study. Experimentally re-infected or vaccinated animals can shed SARS-CoV-2 through upper respiratory tract.^19,26,27,28,29^ In case of Delta AY.1, sgRNA clearance was observed in ¾ animals by 7 DPI but the average viral load remained comparable to that of the primary infection.

Although the risk of re-infection appears to be low in humans, there are reports of varying disease severity in re-infected individuals.^9,11,15,16^ Wang et al, 2021 has reported 68.8%, 18.8% and 12.5% of similar, worse and mild disease severity in re-infection cases.^11^ Severe disease has also been reported with Delta variant re-infection.^15,16^ We did not observe any aggravation of lung disease in the re-infected animals with any of the variants. The lung histopathology score of Delta AY. 1 variant re-infected group showed pneumonic changes of moderate severity in contrast to the mild disease seen in the primary infection group on 7 DPI. When these observations were compared to the overall histopathology score on 3, 7 and 14 DPI after primary infection with Delta AY.1 variant, the severity was comparable. Another indicator of disease severity in hamster model is the body weight loss which was also found minimal here.

Many cytokines has been reported to be increased in severe COVID-19 patients and few like IL-6, IL-8, IL-10 and TNF-α are considered as indicators of severe disease.^30,31^ The increased production of the cytokines can lead to cytokine storm and worsening of the disease prognosis.^32^ Here we have observed increased IL-6 cytokine levels after primary infection in hamsters with SARS-CoV-2 variants. We found no aggravation in cytokine responses post re-infection. IL-6, IL1beta, and TNF increase has been reported in hamsters infected with SARS-CoV-2.^33^ IL-6 is an important cytokine in host responses against viral infection.^33^ The increase observed here could have contributed to the host response in control of infection. Other inflammatory cytokines like IL-4, IFN-Gamma, TNF-alpha also showed slight increase in comparison to control animals during the acute phase of infection. IL-6 level increase was found independent of the lung histopathological score. B.1 variant infected animals showed highest average lung sgRNA load on 7 and 14 DPI and showed highest serum IL-6 levels.

Delta AY.1 variant infected Syrian hamsters produced mild disease in Syrian hamsters characterized by negligible weight loss, a comparative lesser viral load in upper and lower respiratory tract and mild pneumonic changes than Delta variant. Lesser amount of virus shedding in case of Delta AY.1 variant indicates a lower transmission potential of the variant, which needs to be proved by transmission experiments. The lower prevalence rates of the variant i.e., less than 0.5% till date after the initial detection also points to the same^1^. Delta AY.1 variant showed comparable neutralization efficiency against Delta, B.1 and Beta variant. In contrast to this, Delta and B.1 infected animal sera showed a significant lesser neutralization against B.1.351 variant. The IgG response post infection was also highest in case of Delta AY.1. The neutralizing properties of Delta and B.1 variant infected animal sera were also comparable against Delta and Delta AY.1; and B.1 and Delta AY.1 respectively. This indicates that presence of K417N mutation may not confer advantage of immune evasion at least against the Delta and B.1 variants. Yadav et al., 2021 has reported comparable neutralization for the Delta AY.1 and Delta with the sera of naive BBV152 vaccinees, recovered cases with full vaccination and breakthrough cases in comparison to B.1 variant.^35^ Although the disease induced by SARS-CoV-2 in hamsters resemble with humans, many immunological parameters in hamsters are yet to be defined. Our study had limitation of small sample size too.

In conclusion, the present study shows re-infection with Delta variants or B. 1 variant post 3 months of prior B.1 variant infection could reduce the SARS-CoV-2 virus shedding and reduce disease severity. Delta AY.1 variant produced mild disease and generated cross neutralising antibodies against B.1, Delta and Beta variants. The findings indicate that primary infection can produce protective immunity against a further infection for at least 3 months and can reduce severity of infection in hamsters.

## Contributors

PD Yadav and S Mohandas conceived and designed the study. PD Yadav supervised the study. S Mohandas performed the study with the help of PD Yadav and A Shete. PD Yadav, and S Mohandas analysed and interpreted the results. D Nyayanit performed the statistical analysis. R Jain performed the ELISA based experiments. G Sapkal performed the PRNT assays. C Mote performed the histopathology processing.

## Acknowledgments

This study was supported by Indian Council of Medical Research as an intramural grant to ICMR-National Institute of Virology, Pune. Authors acknowledge the laboratory support received from Mr Manoj Kadam, Mr Abhimanyu Kumar, Mr Annasaheb Suryawanshi, Ms. Pranita Gawande, Mrs Ashwini Waghmare, Mrs Kaumudi Kalele, Mr Prasad Sarkale, Mr Shreekant Baradkar, staff of ICMR-National Institute of Virology Pune.

## Data Sharing Statement

All the data pertaining to the study are available in the manuscript or in the supplementary materials.

---

**Supplementary Figure 1:**
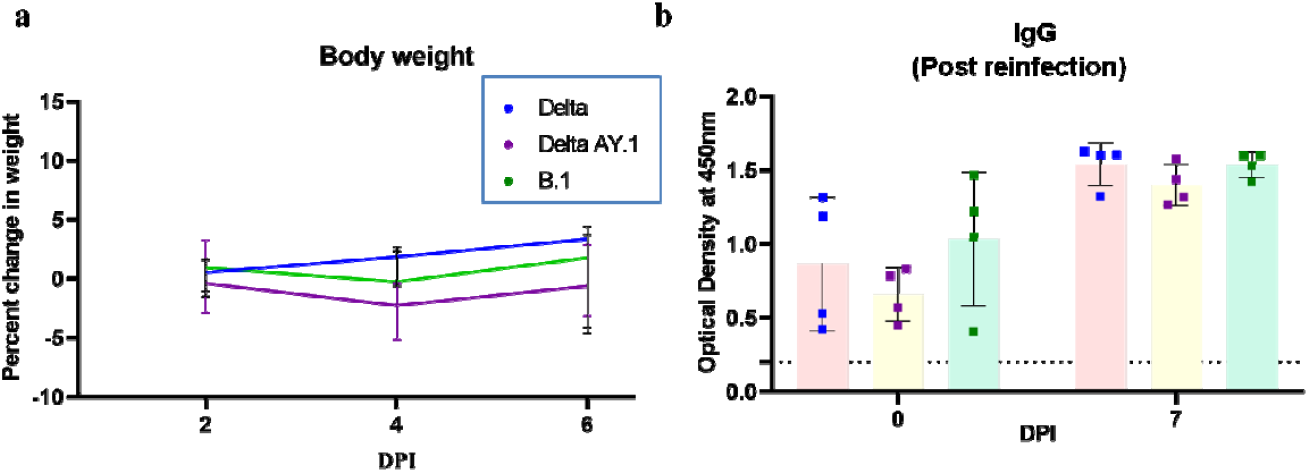
Body weight changes and IgG response in hamsters post re-infection. a) Line graph representing percent body weight change in hamsters on 2, 4 and 6 DPI. b) Scatter plot depicting Optical Density measured by ELISA in hamster sera on day 0 and 7 post re-infection. Limit of detection is assay is depicted as the dotted line.

